# Mapping scientific cosmopolitanism: scientists with publication records substantially split across multiple countries

**DOI:** 10.64898/2026.10.01.755970

**Authors:** John P.A. Ioannidis, Jeroen Baas

**Author notes:** Correspondence: John P.A. Ioannidis, MD, DSc, 3180 Porter Drive, Room A117, Stanford Research Park, Palo Alto, CA 94304, USA.

## Abstract

We used Scopus data on 243,949 top-cited scientists to map scientific cosmopolitanism, defined as having at least two different countries that contribute alone at least 10% of an author’s published record as indexed in Scopus. 49,707 (20.38%) of the top-cited authors were cosmopolitan based on this definition. Based on current author affiliation, rates varied widely (9-76%) across countries with at least 250 top-cited authors with lowest rates in Japan (9%), USA (12%) and Iran (13%). Countries with few top-cited authors had almost always high rates of cosmopolitanism. The proportion of top-cited authors who are cosmopolitan has steadily increased over time in younger generations among those who started publishing after 1965 reaching 43% recently. Large differences were also seen across scientific subfields (1% in General Psychology & Cognitive Sciences to 46% in Econometrics). Among the 100 pairs of countries that were the most commonly expected, the observed to expected ratio ranged from 6.44 for Switzerland-Germany to 0.13 for Italy-Japan. Across the 30 country pairs that were most over-represented, countries shared borders, languages, most common religion pairs, or any of these features in 15, 19, 26, and 29 cases, respectively. The map of cosmopolitan authors reveals a large and increasing proportion of the most influential scientists who share two or more scientific homes that each has made a substantial contribution to their career. Cosmopolitanism among scientists requires better study to help facilitate its potential positive contributions not only to individual scientists but to science and society more broadly.

---

The scientific community transcends national borders, and science is a universal enterprise. Substantial attention has been given to international collaboration and teamwork between scientists from different countries (1–3). However, the extent to which single scientists materialize their career across different countries is less well studied. Mobility of scientists across borders is influenced by multiple factors (4–8) that create unfavorable circumstances for science in some countries (and hence brain drain) and more favorable circumstances in others (and hence brain gain). Moreover, available opportunities for and obstacles to scientific migration may change. Among scientists who migrate across different countries, most eventually operationalize their productive scientific career in only one country. E.g. they may complete basic training in one country and then migrate to another; then all or almost all their publications may originate from one country. However, for some scientists, substantial portions of their published work may come from different countries. E.g. such scientists may already have a substantial published corpus when they migrate; and/or they may return to their country of origin and demonstrate fruitful publication records from both countries. Some other scientists may work in multiple countries during their career.

Here, we aimed to map scientific cosmopolitanism among a comprehensive set of top-cited scientists with the highest citation impact across science. We approached scientific cosmopolitanism using an operational threshold of having at least two different countries that contribute alone at least 10% of an author’s published record as indexed in Scopus. We used science-wide data to understand how common this pattern is, whether it involves some countries and some scientific fields and subfields more than others, whether its prevalence has changed over time, and whether specific countries are highly likely or unlikely to co-exist in scientists’ careers.

## METHODS

### Database of evaluated top-cited authors

We used the August 2026 iteration of the database of top-cited scientists (9) which is annually updated based on Scopus data (10) and captures publications and citations up to end of 2025. We used data on career-long impact that select scientists for inclusion based on a composite score. The composite is computed from 6 indicators (11–13) including number of citations, h-index, co-authorship-adjusted hm index, number of citations as single author, number of citations as single or first author, and number of citations as single, first, or last author. Scientists at the top-2% of the composite score in each of 174 scientific disciplines (classified per Science Metrix (14) and those among the top-100,000 of the composite score across all science are included.

### Definition of cosmopolitan scientists

For each scientist, we used Scopus data to map the countries with which they were affiliated across the totality of their published works indexed in Scopus; and for each country, we noted how many published works carried an affiliation in that country. The attribution of a link between authors’ indicated affiliation on a paper contains a country assignment, which was further enriched to recognize additional countries for cases the single text referred to multiple affiliations with different countries. This step also normalizes the existing country assignments of the publications, by preferring the country or region of the institution as was applied in a previous study (. We noted that it was common for scientists in general (and even more for those with many published works) to have single or few published works coming from a country that was not their most frequent country. However, a large share such sporadic occurrences were artefacts of Scopus coding, e.g. when many countries and authors were listed without clear specific author-country pairing in some large collaborative project or guideline; or may have represented genuine provenance of some early work of a scientist from a country of origin before migrating to the country where they ran the bulk of their career. To avoid artefacts and situations where one country was overwhelmingly responsible for one’s career, we defined cosmopolitan scientists as those who had at least two countries linked to at least h=10% of their career-long published work. This threshold ensured that different countries could each claim a considerable portion of the scientist’s work. Each author-paper record was classified as having no country, one or two usable countries, or three or more countries. Only records with one or two usable countries were used in the country counts. Records with three or more countries were excluded from the denominator because of their higher risk of introducing ambiguous author-country links. Given the inclusion of author-paper records with two usable countries, the sum of h values for an author across all the countries could have exceeded 100%. Countries with h<10% are not counted as contributing to the cosmopolitanism of an author.

Of note, we use the term “country” in a broad sense based on the taxonomy of Scopus which covers regions beyond United Nations member states. The reason is that the Scopus taxonomy covers several territories, islands, and autonomous regions in addition to the member states. The data extract of this study covers 184 distinct eligible regions that had at least one top-cited scientist.

We derived a random sample of 100 authors from top-cited authors with cosmopolitan behavior, and we examined for each of them in depth their Scopus records and online available information about their CV and biography to see if for any of them the cosmopolitan classification was an artefact.

### Analyses

We present descriptive statistics that map scientific cosmopolitanism among top-cited authors. Specifically, we show how many top-cited authors had 2,3,4,5,6 or 7 eligible countries using the 10% threshold; and the proportion of top-cited authors who are also cosmopolitan across different countries, fields, and subfields per Science Metrix, publication age (years since first publication), and number of total published items. Multivariable regression explore the independent associations of these factors with cosmopolitanism.

Finally, we evaluate whether specific countries co-exist very frequently or very infrequently in pairs in the same top-cited author’s career. For this purpose, we calculate the observed-to-expected (O/E) ratio of pairing of countries for cosmopolitan authors. The expected number of pairings E for two countries x1 and x2 is obtained by

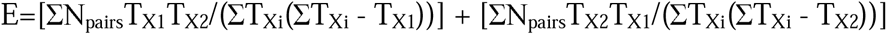

where ΣΝ_pairs_ is the total number of country pairs across all cosmopolitan top-cited authors, T_X1_is the number of top-cited authors whose eligible career-country set contains x1, and T_X2_ similarly the number of top-cited authors whose eligible career-country set contains an x2 address, while ΣT_Xi_ denotes the sum of these country-incidence counts over all eligible countries. Of note, in the total data, we had ΣΝ_pairs_=69,481 pair events.

We also hypothesized that pairing of countries (or lack thereof) in the careers of cosmopolitan authors may be affected by proximity and shared language and culture/religion. Information on whether countries share borders (land or at least sea), official languages (considering de facto language(s) when there was no official one), and most common religion (excluding people who adhere to no religion) was compiled from online searches with the help of Google AI. We also used the EF English Proficiency Index to identify countries with very high English proficiency in the adult population besides those where English is the official or de facto language (15) (https://en.wikipedia.org/wiki/EF_English_Proficiency_Index).

## RESULTS

### Overall numbers of cosmopolitan authors

The sum of the published items of the 243,949 top-cited authors was 52,701,923 author-paper records (covering 34,248,493 distinct publications); of those 48,518,607 listed the top-cited author with one country affiliation, 1,617,922 with two, 113,994 with 3 or more, and 2,451,400 did not have a listed country affiliation for the top-cited author. We excluded from further analysis records with 3 or more usable countries, as well as records with no usable countries, because they may have high frequency of artefacts. Of the 243,949 top-cited authors, 49,707 (20.38%) were cosmopolitan based on our definition. This included 41,760 top-cited authors with substantial publication record from 2 countries, 6,847 from 3 countries, 969 from 4 countries, 121 from 5 countries, 9 from 6 countries, and 1 from 7 countries.

### Validation data

In a random sample of 100 top-cited authors who were classified as being cosmopolitan, for 4 of them, the cosmopolitan profile was an artefact. This included a scientist who had his whole career in Serbia (initially as Yugoslavia, then Sebia), but was listed as having been in Serbia and Croatia; two scientists who worked in the USA but were classified as having been in USA and Finland (because of their affiliation with Bell Labs which were then acquired by Nokia), and one scientist who worked in France but was classified as having been in both France and Switzerland (albeit he had extensive collaborations in Switzerland). The other 96 top-cited authors indeed had spent substantial time in their careers in two or more countries and had published a substantial share of their work from 2 or more countries. Therefore, the false positive rate was 4% (95% Clopper-Pearson interval 1.1-9.9%).

### Cosmopolitan authors per country

Table 1 shows the countries that had at least 250 top-cited authors, in declining order of their proportion of cosmopolitan authors. As shown, the vast majority (76%) of top-cited authors from United Arab Emirates and most (58%) top-cited authors from Saudi Arabia were cosmopolitan and proportions exceeding 40% were seen also in Hong Kong, Singapore, Ireland, Switzerland, and Egypt. The lowest proportions were seen in Japan (9%), USA (12%), Iran (13%), and Italy (16%), while Portugal, Brazil, Taiwan, India and France also had proportions below the global average. Except for Ireland, Greece and Hungary, all EU countries had proportions less than 25%.

**Table 1.** Proportion of cosmopolitan authors according to their current region (limited to regions with at least 250 top-cited authors)

| **Current region** | **Total top-cited authors** | **Cosmopolitan, N** | **Percentage** |
| --- | --- | --- | --- |
| United Arab Emirates | 470 | 355 | 75.53% |
| Saudi Arabia | 1071 | 621 | 57.98% |
| Singapore | 1154 | 557 | 48.27% |
| Hong Kong | 1619 | 771 | 47.62% |
| Ireland | 963 | 453 | 47.04% |
| Switzerland | 3759 | 1658 | 44.11% |
| Egypt | 676 | 288 | 42.60% |
| Pakistan | 321 | 123 | 38.32% |
| Chile | 251 | 95 | 37.85% |
| Austria | 1566 | 577 | 36.85% |
| New Zealand | 1224 | 447 | 36.52% |
| Australia | 8551 | 2876 | 33.63% |
| South Africa | 963 | 317 | 32.92% |
| Israel | 2293 | 751 | 32.75% |
| Greece | 1243 | 379 | 30.49% |
| Norway | 1557 | 470 | 30.19% |
| Hungary | 441 | 131 | 29.71% |
| Malaysia | 515 | 149 | 28.93% |
| Canada | 10063 | 2860 | 28.42% |
| Mexico | 461 | 124 | 26.90% |
| Russia | 1133 | 301 | 26.57% |
| Thailand | 343 | 90 | 26.24% |
| China | 14582 | 3630 | 24.89% |
| Denmark | 2309 | 566 | 24.51% |
| Sweden | 3661 | 891 | 24.34% |
| Argentina | 279 | 67 | 24.01% |
| Romania | 253 | 60 | 23.72% |
| South Korea | 2889 | 674 | 23.33% |
| Czechia | 622 | 145 | 23.31% |
| Germany | 12809 | 2974 | 23.22% |
| United Kingdom | 21351 | 4891 | 22.91% |
| Belgium | 2294 | 510 | 22.23% |
| Spain | 4103 | 905 | 22.06% |
| Netherlands | 5001 | 1057 | 21.14% |
| Turkey | 1479 | 306 | 20.69% |
| Poland | 1581 | 327 | 20.68% |
| Finland | 1612 | 332 | 20.60% |
| India | 3898 | 747 | 19.16% |
| France | 7613 | 1442 | 18.94% |
| Portugal | 787 | 149 | 18.93% |
| Taiwan | 2025 | 365 | 18.02% |
| Brazil | 1304 | 218 | 16.72% |
| Italy | 7363 | 1147 | 15.58% |
| Iran | 1380 | 180 | 13.04% |
| United States | 90921 | 11041 | 12.14% |
| Japan | 9097 | 814 | 8.95% |

Across all countries with fewer than 250 top-cited authors (see Supplement S1), 1,827 of the 3,851 top-cited authors were cosmopolitan, i.e. 47.44%. Of the 105 countries with at least 2 top-cited authors, 102/105 had a higher proportion of cosmopolitan authors than the global 20.38% rate (exceptions were Slovenia, Liberia and Montenegro). In the 25 countries with only 1 top-cited author, 14/25 of these authors were cosmopolitan.

### According to publication age

As shown in Figure 1, cosmopolitanism was relatively common in the early days of science (69/306, 22.55% among top-cited authors who first published before 1930 for Scopus-indexed work) and its frequency declined among scientists who started publishing in 1930-1959 (998/6,768, 14.75%). It reached a low point among those who started publishing in 1960-1964 (789/5,827, 13.59%). Since then, it has become increasingly more frequent among scientists starting to publish in more recent years, exceeding 40% (40.13%, 2,707/6,746) for those starting to publish in 2010-2024, and further escalating to 42.70% (678/1,588) for those starting to publish in 2015-2019, and 42.86% (33/77) for those starting to publish in 2020-2024.

**Figure 1.**
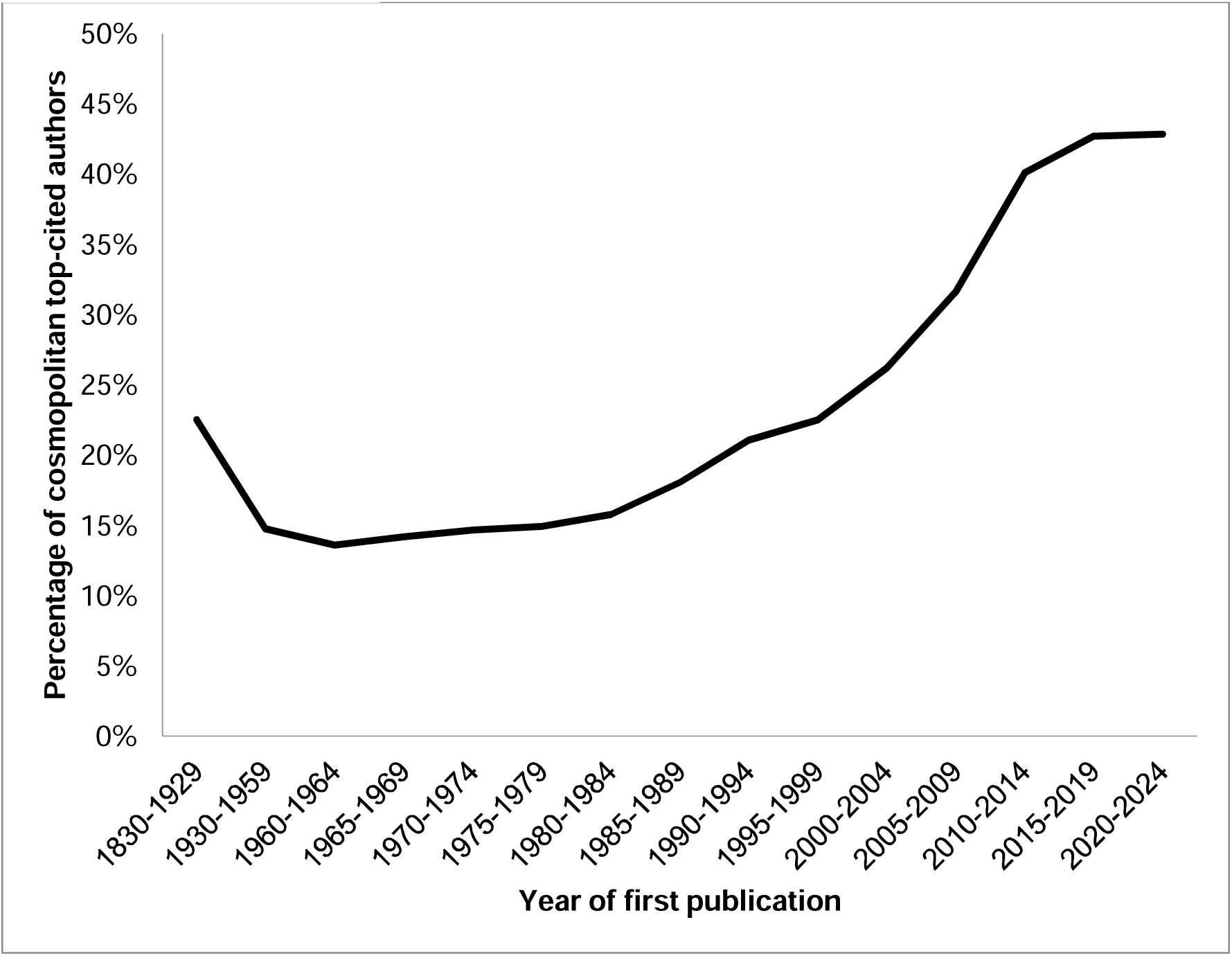
Proportion of top-cited authors who are cosmopolitan according to the year of their first publication

### According to scientific discipline

Based on the 20 Science Metrix fields classification, the highest proportions of cosmopolitan top-cited authors were seen in Economics & Business (1,489/4,838, 30.78%), followed by Physics & Astronomy (6,526/21,480, 30.38%). Conversely, the lowest were seen in Public Health & Health Services (586/4,440, 13.20%), followed by Clinical Medicine (11,001/74,888 = 14.69%). Ten of the 174 subfields had proportions higher than 35% (Econometrics was the highest at 46.05%, followed in decreasing order by Nuclear & Particle Physics, Economic Theory, Sport, Leisure & Tourism, Architecture, Mathematical Physics, Computation Theory & Mathematics, General Physics, Tropical Medicine, and Nanoscience & Nanotechnology). Thirteen subfields had proportions lower than 10% (General Psychology & Cognitive Sciences was the lowest at 1.16%, followed in increasing order by Horticulture, Psychoanalysis, Family Studies, Otorhinolaryngology, Allergy, General Clinical Medicine, Orthopedics, Social Work, Substance Abuse, Gerontology, Pediatrics, and Dermatology & Venereal Diseases). Full data per field and subfield appear in the Supplement S2.

### According to number of papers published in career

The number of published items in a top-cited author’s career was inversely related to the probability of being cosmopolitan (decrease from 13,616/56,617 [24.05%] for those with 0-100 published items, to 18,801/89,402 [21.03%] for those with 101-200, 12,866/71,537 [17.99%] for those with 201-400, and 4,424/26,393 [16.76%] for those with 401 or more).

### Multivariable regression

Table 2 shows the results for multivariable regression for the outcome of being a cosmopolitan author. The patterns seen in univariable analyses were largely retained regarding scientific fields, countries, number of publications, and year of first publication. After adjusting for scientific field and country, cosmopolitanism increased in more recent years and among top-cited authors with fewer publications.

**Table 2.** Multivariable model for cosmopolitan top-cited authors.

| **FACTORS** | **Odds ratio** | **Lower 95% CI** | **Upper 95% CI** |
| --- | --- | --- | --- |
| **Scientific field** |  |  |  |
| Clinical Medicine | Reference |  |  |
| Agriculture, Fisheries & Forestry | 0.83 | 0.78 | 0.89 |
| Biology | 1.35 | 1.28 | 1.43 |
| Biomedical Research | 1.49 | 1.43 | 1.56 |
| Built Environment & Design | 1.10 | 0.97 | 1.24 |
| Chemistry | 1.21 | 1.16 | 1.27 |
| Communication & Textual Studies | 1.18 | 1.02 | 1.35 |
| Earth & Environmental Sciences | 1.38 | 1.31 | 1.47 |
| Economics & Business | 1.94 | 1.81 | 2.08 |
| Enabling & Strategic Technologies | 1.29 | 1.24 | 1.35 |
| Engineering | 1.08 | 1.04 | 1.13 |
| Historical Studies | 1.47 | 1.28 | 1.69 |
| Information & Communication Technologies | 1.70 | 1.63 | 1.77 |
| Mathematics & Statistics | 1.99 | 1.82 | 2.18 |
| Philosophy & Theology | 1.66 | 1.36 | 2.02 |
| Physics & Astronomy | 2.62 | 2.53 | 2.72 |
| Psychology & Cognitive Sciences | 1.22 | 1.11 | 1.32 |
| Public Health & Health Services | 0.72 | 0.66 | 0.79 |
| Social Sciences | 1.00 | 0.93 | 1.07 |
| Visual & Performing Arts | 1.13 | 0.69 | 1.85 |
| **Number of publications** |  |  |  |
| Up to 100 | Reference |  |  |
| 101-200 | 0.84 | 0.81 | 0.86 |
| 201-400 | 0.70 | 0.67 | 0.72 |
| 401 or more | 0.66 | 0.63 | 0.69 |
| **Country** |  |  |  |
| USA | Reference |  |  |
| ARE | 17.21 | 13.79 | 21.49 |
| ARG | 2.22 | 1.68 | 2.93 |
| AUS | 3.73 | 3.54 | 3.93 |
| AUT | 4.03 | 3.62 | 4.48 |
| BEL | 2.14 | 1.93 | 2.37 |
| BRA | 1.41 | 1.21 | 1.63 |
| CAN | 3.05 | 2.90 | 3.20 |
| CHE | 5.64 | 5.26 | 6.05 |
| CHL | 3.85 | 2.97 | 4.99 |
| CHN | 1.74 | 1.66 | 1.83 |
| CZE | 1.96 | 1.62 | 2.38 |
| DEU | 2.06 | 1.96 | 2.16 |
| DNK | 2.39 | 2.17 | 2.64 |
| EGY | 3.69 | 3.12 | 4.35 |
| ESP | 1.86 | 1.72 | 2.01 |
| FIN | 1.90 | 1.68 | 2.15 |
| FRA | 1.61 | 1.51 | 1.71 |
| GBR | 2.13 | 2.05 | 2.21 |
| GRC | 2.86 | 2.52 | 3.25 |
| HKG | 6.06 | 5.46 | 6.73 |
| HUN | 3.04 | 2.45 | 3.77 |
| IND | 1.29 | 1.19 | 1.41 |
| IRL | 6.55 | 5.73 | 7.49 |
| IRN | 0.71 | 0.60 | 0.83 |
| ISR | 3.49 | 3.19 | 3.82 |
| ITA | 1.28 | 1.20 | 1.37 |
| JPN | 0.73 | 0.68 | 0.79 |
| KOR | 1.80 | 1.64 | 1.97 |
| MEX | 2.40 | 1.93 | 2.97 |
| MYS | 2.28 | 1.86 | 2.78 |
| NLD | 2.00 | 1.86 | 2.15 |
| NOR | 3.13 | 2.80 | 3.51 |
| NZL | 4.41 | 3.90 | 4.98 |
| PAK | 2.64 | 2.06 | 3.40 |
| POL | 1.51 | 1.33 | 1.71 |
| PRT | 1.49 | 1.24 | 1.78 |
| ROU | 1.69 | 1.25 | 2.28 |
| RUS | 2.14 | 1.87 | 2.46 |
| SAU | 6.75 | 5.91 | 7.71 |
| SGP | 6.03 | 5.34 | 6.82 |
| SWE | 2.43 | 2.24 | 2.63 |
| THA | 2.44 | 1.89 | 3.16 |
| TUR | 1.23 | 1.07 | 1.41 |
| TWN | 1.35 | 1.20 | 1.52 |
| ZAF | 3.52 | 3.06 | 4.06 |
| Other countries (with less than 250 top-cited authors) | 5.26 | 4.90 | 5.65 |
| **Year of first publication** |  |  |  |
| 1960-1964 | Reference |  |  |
| 1830-1929 | 1.72 | 1.30 | 2.29 |
| 1930-1959 | 1.14 | 1.03 | 1.26 |
| 1965-1969 | 1.02 | 0.93 | 1.13 |
| 1970-1974 | 1.09 | 1.00 | 1.19 |
| 1975-1979 | 1.14 | 1.05 | 1.24 |
| 1980-1984 | 1.21 | 1.11 | 1.31 |
| 1985-1989 | 1.36 | 1.25 | 1.48 |
| 1990-1994 | 1.55 | 1.43 | 1.68 |
| 1995-1999 | 1.66 | 1.53 | 1.80 |
| 2000-2004 | 1.98 | 1.82 | 2.15 |
| 2005-2009 | 2.49 | 2.28 | 2.71 |
| 2010-2014 | 3.33 | 3.03 | 3.67 |
| 2015-2019 | 3.11 | 2.71 | 3.57 |
| 2020-2024 | 3.31 | 1.92 | 5.72 |

### Pairing of countries with higher-than-expected and lower-than-expected frequency

Among the 100 pairs of countries that were the most expected (details in Supplement S3), 5 had O/E ratios >2: Switzerland-Germany (6.44); Australia-UK (2.98); Germany-Netherlands (2.76); UK-Singapore (2.18); UK-Netherlands (2.02). Conversely, 5 countries had O/E ratios <0.33: UK-Taiwan (0.32); Japan-Netherlands (0.29); Malaysia-USA (0.24); China-Italy (0.21); Italy-Japan (0.13). In the 5 most over-represented pairs, the paired countries shared borders in 3/5; official (or de facto) languages in 3/5 and in the 5/5 when we also considered very high English proficiency; and most common religion in 4/5. Conversely, among the 5 lowest O/E ratios, countries never shared borders, official or de facto language or very high English proficiency, and most common religion.

Overall, 2,617 different country pairs were observed to have at least 1 top-cited author (details in Supplement S4). This is 15.54% of the theoretically possible 16,836 region pairs across the 184 distinct eligible country codes extracted in the study.

For the 34 countries with at least 800 top-cited authors based on current affiliations, the number of theoretically possible pairs is 561. Almost all (551/561, 98.22%) were observed to have at least 1 top-cited cosmopolitan author. The 10 missing country pairs were Hong Kong and Ireland, Israel and Saudi Arabia, Denmark and Taiwan, India and Greece, New Zealand and Greece, Israel and Turkey, Turkey and Ireland, Taiwan and Greece, Brazil and Greece, Taiwan and Brazil. Countries in these pairs were far apart geographically and never shared official or de facto language or very high English proficiency. For 2/10 pairs, Christianity was the most common religion (although denominations differed). Some pairs had also a history of strained relationships.

A total of 182 pairs of countries had E>0.1 and O/E>10 (see Figure 2 and Supplement S5), i.e. the observed number of cosmopolitan top-cited authors was more than 10 times higher than the expected number. A dominant cluster was comprised of countries in the Arab and/or Muslim world, including prominently Saudi Arabia (n=16 occurrences among these 182 pairs), Malaysia (n=14 occurrences), United Arab Emirates (n=11), Pakistan (n=9), Turkey (n=9), Kazakhstan (n=8), Iran (n=7), Kuwait (n=7), Oman (n=7), Egypt (n=6), and Qatar (n=6) and others with fewer occurrences in over-represented pairs. Additional countries with at least 6 occurrences among these highly over-represented pairs were France (n=11), Vietnam (n=9), South Africa (n=8), Australia (n=7), Spain (n=7), Switzerland (n=6), Cyprus (n=6), UK (n=6), and Russia (n=6). The five highest O/E ratios were United Arab Emirates-Jordan (196.15), Azerbaijan-Turkey (175.57), Iraq-Malaysia (159.73), Egypt-Saudi Arabia (142.45), and Indonesia-Malaysia (116.31).

**Figure 2.**
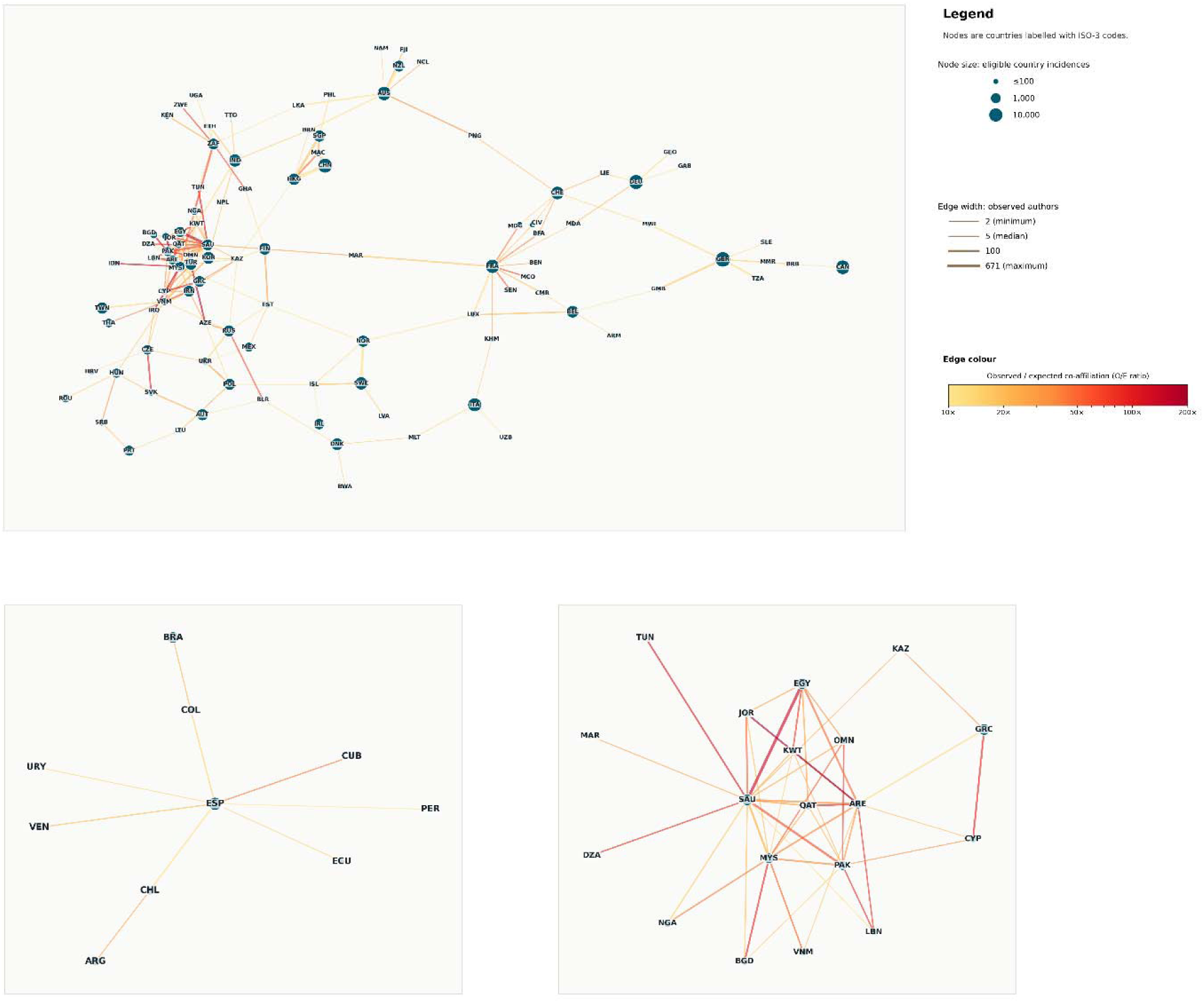
Strongly enriched country and region co-affiliation pairs among top-cited authors: network visualization of the 182 pairs of countries and regions where E>0.1 and O/E>10; main panel A shows 101 nodes, panel B shows 10 nodes with no connection meeting the threshold to nodes in panel A. Panel C shows the detailed view of the one-hop nodes surrounding the ARE-EGY-SAU cluster.

Table 3 shows the 30 country pairs with E>0.1 and the highest O/E ratios. Among them, there were shared borders in 15/30, shared official or de facto languages in 19/26, and shared major religion in 26/30 (most commonly Islam, in 18/30). In 29/30 pairs, the countries shared at least one of these features; the only exception was Iran-Vietnam. Countries with Arab and/or Islam background appeared in 22/30 pairs, while the other 8 highly over-represented pairs reflected diverse situations with strong cultural and/or historical ties (Czech Republic-Slovakia, Greece-Cyprus, France-Monaco, Russia-Belarus, Hong Kong-Macao, France-Madagascar) and the attraction of South Africa for scientists in other African countries (Zimbabwe, Ghana).

**Table 3.** Country pairs with the highest O/E ratio of cosmopolitan top-cited authors than (among those with E>0.1): shared borders, official or de facto languages, and most common religion (among those who have a religion)

| Country 1 | Country 2 | O | E | O/E | Border | Language | Religion |
| --- | --- | --- | --- | --- | --- | --- | --- |
| UAE | Jordan | 30 | 0.15 | 196.1 | No | Yes (Arabic) | Yes (Islam) |
| Azerbaijan | Turkey | 23 | 0.13 | 175.6 | Yes | No (Azerbaijani vs. Turkish) | Yes (Islam) |
| Iraq | Malaysia | 23 | 0.14 | 159.7 | No | No (Arabic vs. Malay) | Yes (Islam) |
| Egypt | Saudi Arabia | 348 | 2.44 | 142.5 | Yes | Yes (Arabic) | Yes (Islam) |
| Indonesia | Malaysia | 16 | 0.14 | 116.3 | Yes | No (Indonesian vs. Malay) | Yes (Islam) |
| Saudi Arabia | Tunisia | 20 | 0.18 | 109.2 | No | Yes (Arabic) | Yes (Islam) |
| Czech Republic | Slovakia | 18 | 0.17 | 103.2 | Yes | No (Czech vs. Slovak) | Yes (Christianity) |
| UAE | Qatar | 21 | 0.22 | 97.3 | Yes | Yes (Arabic) | Yes (Islam) |
| Bangladesh | Malaysia | 25 | 0.27 | 92.6 | No | No (Bengali vs. Malay) | Yes (Islam) |
| Lebanon | Pakistan | 11 | 0.12 | 90.3 | No | No (Arabic vs. Urdu) | Yes (Islam) |
| Cyprus | Greece | 34 | 0.38 | 89.6 | Yes | Yes (Greek) | Yes (Christianity) |
| Oman | Pakistan | 9 | 0.10 | 86.3 | Yes | No (Arabic vs. Urdu) | Yes (Islam) |
| UAE | Lebanon | 10 | 0.12 | 86.0 | No | Yes (Arabic) | Yes (Islam) |
| Algeria | Saudi Arabia | 12 | 0.15 | 80.3 | No | Yes (Arabic) | Yes (Islam) |
| Egypt | Kuwait | 17 | 0.22 | 79.0 | No | Yes (Arabic) | Yes (Islam) |
| South Africa | Zimbabwe | 9 | 0.12 | 74.9 | Yes | Yes (English/Zulu/Xhosa) | Yes (Christianity) |
| France | Monaco | 8 | 0.12 | 68.3 | Yes | Yes (French) | Yes (Christianity) |
| Belarus | Russia | 11 | 0.17 | 66.4 | Yes | Yes (Russian) | Yes (Christianity) |
| Ghana | South Africa | 9 | 0.14 | 64.7 | No | Yes (English) | Yes (Christianity) |
| Nigeria | South Africa | 39 | 0.61 | 64.4 | No | Yes (English) | No (Islam vs. Christianity) |
| Pakistan | Saudi Arabia | 83 | 1.33 | 62.6 | No | No (Urdu vs. Arabic) | Yes (Islam) |
| Iran | Vietnam | 29 | 0.50 | 57.6 | No | No (Persian vs. Vietnamese) | No (Islam vs. Folk) |
| Malaysia | Oman | 9 | 0.16 | 56.0 | No | No (Malay vs. Arabic) | Yes (Islam) |
| Jordan | Saudi Arabia | 25 | 0.46 | 54.0 | Yes | Yes (Arabic) | Yes (Islam) |
| UAE | Kuwait | 6 | 0.11 | 53.8 | Yes | Yes (Arabic) | Yes (Islam) |
| France | Senegal | 14 | 0.26 | 53.1 | No | Yes (French) | No (Christianity vs. Islam) |
| Malaysia | Vietnam | 13 | 0.25 | 52.7 | Yes | No (Malay vs. Vietnamese) | No (Islam vs. Folk) |
| UAE | Egypt | 39 | 0.81 | 48.3 | No | Yes (Arabic) | Yes (Islam) |
| Hong Kong | Macau | 29 | 0.61 | 47.7 | Yes | Yes (Cantonese/Chinese) | Yes (Chinese Folk) |
| France | Madagascar | 8 | 0.18 | 45.5 | Yes | Yes (French) | Yes (Christianity) |

## DISCUSSION

Science-wide mapping shows that one in five top-cited scientists have 2 or more countries as the origin of a substantial share of their published work. The proportion of such cosmopolitan behavior has steadily increased in the last six decades and has exceeded 40% among those who started publishing after 2010. Countries have differed up to more than 8-fold in the proportion of their top-cited authors who are cosmopolitan with very low rates in Japan, USA, and Iran and very high rates in UAE, Saudi Arabia, Singapore, Hong Kong, Ireland, and Switzerland as well as in the vast majority of countries that have few top-cited authors. Up to about 40-fold differences were seen across scientific subfields. Some pairings of countries were very strongly overrepresented (given the expectation from their share of top-cited authors) while others were under-represented and even totally absent. Proximity (shared borders) and cultural factors such as shared (or not) language and religion were prominently associated with this variability.

Scientific cosmopolitanism, as defined here, needs to be seen in the context of the broader forces that shape scientific mobility (including its facilitators and obstacles), brain drain, and brain gain. Countries that have the highest shares of cosmopolitan top-cited authors and have many top-cited authors have made explicit efforts to attract talent from other countries and have seen formidable brain gain (16–18).

In some cases, policies may facilitate spurious allocation of credit to a given institution and thus its country without true scientific migration. For example, some universities in Saudi Arabia have tried to lure highly cited scientists from other countries to put Saudi affiliations in their work for a generous payment (19,20). In another prominent example among the 30 most over-represented country pairs with the highest E/O ratio, Iran-Vietnam was the only one where there were no shared borders, language and religion and the over-representation may be driven by highly cited Iranian scientists using affiliations of Vietnamese institutions without moving to Vietnam. Some Vietnamese institutions may pay highly cited scientists to adopt their affiliation in order to improve their ranking (20) and Iranian scientists may seek a better salary and an affiliation that may be causing less suspicion. Some of the multiple affiliation cases may also be associated with spurious research practices, hyper-prolific behavior (21,22), and higher risk of retractions (23).

However, in most cases, brain gain has been genuine; and attraction of talent has been accompanied by growth in R&D, investment in scientific infrastructure, and overall prosperity driven by science and technological advances. Conversely, for countries with very few top-cited authors, the very high rates of cosmopolitanism may reflect largely genuine brain drain patterns. Talented scientists in these countries work in very difficult conditions and working in a country with better resources may be the best, if not only, way to make substantial progress. Proximity and cultural matching factors may shape much of the migration destinations for these researchers. The extent to which these scientists can eventually offer back to their original country needs better study (24).

Countries with low proportion of cosmopolitan scientists may exhibit different patterns of isolationism, as exemplified by the contrast of Japan, USA, and Iran. Japan (26) may exhibit relatively low scientific migration in terms of both immigration and emigration. The language barrier may diminish its attractiveness for foreign talent and Japanese scientists are also unlikely to have a career abroad given the high standards they can find in their country. USA is far more successful for attracting talent from other countries and this has been a key contributor to the overall success of USA in science and technology (27–30). Current threats to impose barriers for such scientific migration are deemed to threaten USA hegemony in science. However, even though a large share of top-cited scientists currently in the USA are foreign-born and/or trained, typically all or almost all their publication record has been generated from the USA. Lastly, Iran exhibits severe isolationism with very large brain drain (31). Iran-born cosmopolitan top-cited scientists thus often appear under their countries of destination (current address) in our analyses.

Differences in cosmopolitanism across scientific fields and subfields may reflect scientific infrastructure required to do high-impact work and long-standing norms for international collaboration and mobility. Communities such as high energy and particle physics have high mobility related to where experiments can be done. Economics and business fields also share global standards and global markets. Conversely, other fields such as those in biomedicine may have advanced teamwork and collaborative structures (32) but typically researchers do not have to migrate to collaborate with colleagues around the world.

Our map of cosmopolitan publication behavior shows that pairing of countries in authors’ careers is far from random. Over-represented pairs tend to share geographic proximity and cultural features such as language and religion. Besides official and de facto languages, English especially is the connecting lingua franca language for science also for many other countries (33). Cultural values, as proxied by Arabic language and Islam religion, may be instrumental in governing mobility across countries in the Middle East and beyond. Home countries and institutions may occasionally apply pressure to graduates to continue using their original home affiliation (34). Conversely, some country pairs were grossly under-represented or even totally absent and almost always this reflected distance and lack of shared cultural proxies. Our map of country pairings may be compared against maps of scientific collaboration in terms of authorship of papers which also show large heterogeneity (35). However, having co-authors from multiple countries is conceptually different from having authors have their career spent to a substantive degree in multiple countries.

Some limitations of our work should be discussed. First, we focused on the set of top-cited authors, and these represent a small share of the scientific community. However, these authors are the most influential in shaping science. Moreover, for most scientists, their classification as cosmopolitan based on the same 10%-of-publications rule would be precarious, since most have published very few items and artefacts of Scopus coding of country would create false classifications even if just one or two papers had wrongly coded country address. Second, our definition used a 10% threshold for publications. The classification of authors as cosmopolitan would have increased slightly to 22.7% if the threshold was based on the share of citations obtained instead of publications. Specifically, an additional 13,936 top-cited authors would be cosmopolitan based on the citation-based definition, while 8,217 would lose cosmopolitan status when moving from the publication-based to the citation-based definition. However, using a 10% threshold for citations might have increased the impact of Scopus coding artefact, since some extremely highly cited guidelines have ambiguous country coding for authors. Third, the definition of top-cited author and the relevant citation metrics were taken from previous work (11–13), but the exact operationalization of top-cited authors can always be debated. The same applies to how good citation metrics can be for identifying accurately the very best scientists (36). Nevertheless, the set that we examined is clearly a highly influential cohort. Fourth, we did not collect in-depth information on the individualized trajectories of cosmopolitan authors. This would require additional studies, coupled with different methods, e.g. personal interviews. However, we did examine in depth a random sample of 100 authors classified as cosmopolitan by our methods and found a low rate of classification artefacts. Conversely, probably that a much larger share of top-cited scientists did spend some time in countries other than their primary one without necessarily publishing a substantial share of their work from there. Hence, more leniently defined cosmopolitanism may be substantially more frequent.

Allowing these caveats, our mapping of scientific cosmopolitanism shows that a large share of highly influential scientists share two or more scientific homes that each has made a substantial contribution to their career. For the generation of scientists who have started their career recently and are poised to become the most influential ones in their discipline, these cosmopolitan scientists may be the absolute majority. Influential scientists who train, live and work only in one country throughout their careers may become a minority. Conversely, initiatives that favor isolationism and pose obstacles to scientific migration may halt this phenomenon and this may have negative consequences. Cosmopolitanism among scientists may have positive repercussion with broadening of minds, enhancement of global networking and collaboration, and bridging of differences across countries and nations.

## Notes

### Competing Interest Statement

Jeroen Baas in an employee of Elsevier that produces Scopus

